# The *t* haplotype, a selfish genetic element, manipulates migration propensity in free-living wild house mice *Mus musculus domesticus*

**DOI:** 10.1101/271247

**Authors:** Jan-Niklas Runge, Anna K. Lindholm

## Abstract

Life is built on cooperation between genes, which makes it vulnerable to parasitism. However, selfish genetic elements that exploit this cooperation can achieve large fitness gains by increasing their transmission unfairly relative to the rest of the genome. This leads to counter-adaptations that generate unique selection pressures on the selfish genetic element. This arms race is similar to host-parasite co-evolution. Some multi-host parasites alter the host’s behaviour to increase the chance of transmission to the next host. Here we ask if, similarly to these parasites, a selfish genetic element in house mice, the *t* haplotype, also manipulates host behaviour, specifically the host’s migration propensity. Variants of the *t* that manipulate migration propensity could increase in fitness in a meta-population. We show that juvenile mice carrying the *t* haplotype were more likely to emigrate from and were more often found as migrants within a long-term free-living house mouse population. This result may have applied relevance as the *t* has been proposed as a basis for artificial gene drive systems for use in population control.

## Introduction

The genes within a genome must work together to produce a viable organism, but their interests are not identical [1]. This causes conflict, because not all genes in an organism will be transmitted in equal numbers to the next generation. Consequently, a fair chance of transmission is necessary for cooperation within the genome over evolutionary time. Genes that violate this rule by unfairly increasing their chance of transmission can gain large fitness advantages at the detriment of those that act fairly [2]. This leads to selection for selfish adaptations and, as a result, counter-adaptations to this selfishness, initiating an arms race between these selfish genetic elements and the rest of the genome. This arms race is similar to the one between hosts and parasites, where some parasites even manipulate their hosts. For example, rats infected with the multi-host parasite *Toxoplasma gondii* show decreased fear of cat odour [3]. This is expected to increase the risk of predation by cats, the final host of *T. gondii*, thereby increasing the transmission of the parasite. The parasite furthermore increases the attractiveness of its host, circumventing the avoidance of the infected individual by other rats [3]. Similar manipulations have been observed, for example, in fungi-infected ants that climb vegetation and remain latched onto it post-mortem, which leaves their infested bodies conspicuous to predators [4].

Host defences against parasites and “parasitic” [5,6] selfish genetic elements range from behavioural changes to increased resistance in infected populations. For example, populations of the amphipod *Gammarus pulex* that are not naturally infected with the parasite *Pomphorhynchus laevis* are more sensitive to the parasite’s manipulation than naturally infected populations [7]. This is evidence of an arms race. A similar counter-adaptation to selfish genetic elements is the suppression of the mechanism of the drive. For example, in systems with X chromosome drive in *Drosophila*, which lead to the killing of Y-carrying sperm, some (Y) chromosomes suppress the drive, restoring production of sons [8–11]. Behavioural adaptations are also evident, especially in mating preferences that reduce transmission of parasites or selfish genetic elements. In the woodlouse *Armadillidium vulgare*, males discriminate against “neo-females” infected with feminizing *Wolbachia* bacteria, another type of selfish genetic element[12]. Similarly, females discriminate against individuals carrying a selfish genetic element in stalk-eyed flies [13]. Furthermore, females evolve higher remating rates in response to the presence of a selfish genetic element in *Drosophila pseudoobscura*, which reduces its fitness [14].

Male meiotic drivers are selfish genetic elements that manipulate spermatogenesis to favour the sperm that carry them by harming the sperm that do not [15,16]. This is expected to decrease the competitiveness of a male carrying the meiotic driver by decreasing the number of viable sperm and potentially damaging the driver-carrying sperm as a by-product [16,17]. In consequence, driver-carrying individuals will perform worse [18,19] in sperm competition, in which sperm of different males compete over fertilization. Potentially, the driver carriers might not sire a single offspring despite mating [17] and the driver could go locally extinct [20]. Because of this strong disadvantage, females can be selected to increase sperm competition to decrease the risk of transmitting a driver to their offspring [14,21,22]. In response, the driver could manipulate the male host’s reproductive behaviour as may be the case in *Wolbachia*-infected *Drosophila* that show higher mating rates [23]. Not much is otherwise known about how male meiotic drivers respond to this counter-adaptation that increases the risk of their extinction.

The *t* haplotype (*t*) is a male meiotic driver in the house mouse *Mus musculus*. It consists of a set of genes, making up about 1.5% or 40 Mb of the mouse genome, that are linked by inversions [2,24] and distort Mendelian inheritance patterns so that 90 - 99% of the offspring inherit the *t* from a heterozygous sire [25,26]. It harms its host in at least two ways. The *t* carries recessive lethal alleles, so that *t*/*t* die prenatally [17,27]. In addition, *t* heterozygous (+/*t*) males are very poor sperm competitors, siring only 11%-24% of offspring when mating with a female who also mates with a wildtype male in the same oestrus cycle [17,28]. In house mice, sperm competition intensity varies between populations [29] and is higher in larger populations [30], so that fitness losses of +/*t* males from sperm competition are likely to vary with population demography. This is consistent with a negative association between population size and *t* frequency found in a trapping study [31]. In an intensively monitored free-living large house mouse population, the frequency of the *t* decreased significantly over 6 years until no +/*t* were left [20] while population size increased [32]. Experimental evidence shows that *t* frequency decline in this population is not linked to mate choice against the *t* haplotype [33,34] as found by Lenington et al.[35] in another population, but is influenced by sperm competition [17,20].

The decline of the *t* in the population was even more rapid than a model based on sperm competition predicted [20]. One additional contributing factor could be that +/*t* individuals are more likely to emigrate from the population than +/+. We will use the term ‘emigration’ when we mean leaving the natal population (the first step of dispersal [36]), ‘migration’ when we mean leaving and entering another deme or population [37], and ‘dispersal’ when we mean migrating and then breeding. Early theoretical work predicted that increased dispersal rates should be beneficial for the *t* haplotype by preventing it from extinction due to drift and allowing it to increase in frequency rapidly when dispersing to a suitable population [38]. In this view, a suitable population would be one that has no +/*t* in it, because the fitness of the *t* is frequency dependent, with lower fitness at high *t* frequencies [39]. This is due to negative fitness effects (up to homozygous lethality) of deleterious mutations on the *t* [25]. Combined with the more recent discovery of low sperm competitiveness, the most suitable population for the *t* would therefore be one with as few +/*t* and as little sperm competition as possible, which is expected in smaller populations [30]. A *t* variant that is more likely to disperse to such a population should therefore be at a selective advantage compared to other variants. However, the competitive disadvantage of +/*t* male sperm is also frequency dependant as sperm competition between two +/*t* males does not disfavour the *t* [33]. Considering the negative fitness consequences for *t* under high *t* or high female multiple mating frequencies, populations with both (high *t* frequency and high female multiple mating frequency) are probably rare.

We hypothesized that a *t* mutant that increases the migration propensity of its host generally would more often disperse to suitable populations and would thereby be selected. The increase in migration propensity could be general or be a function of population density (i.e. +/*t* might only emigrate more than +/+ in dense populations where sperm competition is more common [29,30]). This has not been tested yet, but for parasites, theoretical work has demonstrated that they would benefit from manipulating their host’s migration propensity because the parasite’s fitness is influenced by migratory movements of the host [40,41]. However, while there is evidence that parasites can adapt to use host migratory movements for their own advantage [42] and manipulate locomotory behaviours [4], to our knowledge no influence on host migration behaviour has been shown so far. This is probably partly due to how difficult it is to measure migration and dispersal of infected *vs*. non-infected individuals. We analysed juvenile migration from and within an open population of wild house mice (the same as analysed for *t* frequency dynamics by Manser et al.[20]) to investigate if +/*t* individuals are more likely to emigrate than +/+. We found that +/*t* juveniles were more likely to emigrate from the population than +/+, particularly when juvenile densities were high. To our knowledge this is the first evidence of increased migration propensity of carriers of any selfish genetic element in a free-living population. Our research is particularly timely, as the *t* haplotype is proposed as a basis for artificial gene drive systems to eradicate house mouse populations [43,44] and behavioural differences in migration propensity between +/*t* and +/+ would need to be considered in modelling and implementing such systems.

## Results

### Emigration out of the population

We analysed long-term data from a closely monitored population of house mice (*Mus musculus domesticus*) that live in a barn, which they can enter and leave freely (for details see Methods). We defined juvenile emigrants as individuals that disappeared from the population between late pup stage and adulthood (meaning they were not found as adults or as corpses). Based on this, 56% of all individuals born (*N* = 2938) in the years of this analysis who were alive shortly before weaning emigrated. We used a generalized linear mixed modelling approach to investigate the effect of genotype on juvenile emigration propensity, incorporating the covariates of sex, season (main breeding season *vs*. off-season), and juvenile and adult population sizes as well as the year of birth as a random effect.

The most informative model included the genotype and an interaction between the genotype and the juvenile population size (model 2, see Table 1 and the S1 Table). We chose this model to look at effect sizes (Figure 1). This model indicated that +/*t* were more likely to emigrate, particularly with increasing numbers of juveniles in the population (Figure 2). At mean juvenile densities, the probability that a +/*t* juvenile emigrates was 47.5% higher than the probability for a +/+ juvenile (based on model predictions used for Figure 2). A standard deviation increase in juvenile population size increased this difference by 13.3 percentage points. As can be seen in Figure 2, +/*t* and +/+ were similar in their emigration propensity when there were few juveniles in the population, but then diverged with increasing juvenile density. Emigration probability decreased with increasing adult population sizes, but was not differently affected in +/+ and +/*t*. Similarly, being born in the main breeding season and being female increased the probability of emigration for both genotypes.

To test possible alternative explanations for the emigration propensity of +/*t* (like a mortality or condition bias), we analysed data on dead juveniles found in the same time frame. We found no difference between +/+ and +/*t* in the number of dead juveniles *vs*. the number of individuals that did not die or emigrate as juveniles (+/*t*: 17.8% of 90 non-emigrants died as juveniles, +/+: 14.2% of 1424, χ^2^ = 0.62, *p* = 0.43). We decided not to conduct a more detailed model for this comparison because of the limited amount of juvenile +/*t* corpses found (*N* = 20). To ease comparison of this simple mortality analysis with the emigration model, we used the same simple statistical test for the emigration data and again found the difference between +/*t* and +/+ (71.6% of 261 +/*t* that did not die as juveniles emigrated as juveniles, +/+: 54.4% of 2677, χ^2^ = 28.16, *p* = 1.1*e* − 7). We also tested whether there were any differences in the individual body mass as a pup (as a measure of the condition of the pup) between +/+ and +/*t*. We found that +/*t* pups were slightly heavier than +/+ pups (*β* = 0.17*g*, *p* = 0.03, intercept = 6.46g, details in S2 Table), but did not find that the body mass as a pup affects emigration, neither when the genotype was in the model nor when it was absent (models 7 & 8, see Table 1). Thus, we concluded that differences in juvenile emigration between the genotypes cannot be explained by differences in juvenile mortality or condition.

**Table 1:** Overview of models of juvenile emigration out of the study population. Asterisks in model terms indicate interactions. Comparison shows against which other model the model in the row was evaluated. *LRT* indicates the likelihood ratio test statistic of the observed dataset. The *p*-value is the fraction of simulated datasets with *LRT* larger than the observed *LRT* (see Methods). Runs indicate the absolute values on which the *p* is based. The Δ*AIC* is given for comparison with other statistical approaches. The star indicates that these models were restricted by removing individuals without data on pup body mass (see Methods).

| Models | Formula | Comparison | $LRT$ | $p$ | Runs | $\Delta AIC$ |
| --- | --- | --- | --- | --- | --- | --- |
| Null model with covariates | ∼ juvenile population size<br>$+$ juvenile population size<sup>2</sup><br>$+$ adult population size<br>$+$ adult population size<sup>2</sup><br>$+$ season $+$ sex<br>$+$ age when sampled | NA | NA | NA | NA | NA |
| Model 1 | ∼ genotype<br>$+$ null model variables | Null model | 16.00 | 0.0003 | 1/5869 | -14.0 |
| Model 2 | ∼ genotype * juv. pop. size<br>$+$ genotype * juv. pop. size<sup>2</sup><br>$+$ model 1 variables | Model 1 | 11.62 | 0.005 | 26/5815 | -7.62 |
| Model 3 | ∼ genotype * ad. pop. size<br>$+$ genotype * ad. pop. size<sup>2</sup><br>$+$ model 1 variables | Model 1 | 4.24 | 0.10 | 884/8512 | -0.24 |
| Model 4 | ∼ genotype * ad. pop. size<br>$+$ genotype * ad. pop. size<sup>2</sup><br>$+$ model 2 variables | Model 2 | 0.79 | 0.70 | 5402/7681 | +3.21 |
| Model 5 | ∼ genotype * season<br>$+$ model 2 variables | Model 2 | 0.03 | 0.96 | 7092/7355 | +1.97 |
| Model 6 | ∼ genotype * sex<br>$+$ model 2 variables | Model 2 | 1.12 | 0.41 | 2807/6912 | +0.88 |
| Model 7 | ∼ pup body mass<br>$+$ null model variables | Null model* | 0.97 | 0.38 | 3160/8359 | +1.03 |
| Model 8 | ∼ pup body mass<br>$+$ model 2 variables | Model 2* | 1.34 | 0.46 | 3770/8150 | +0.66 |

**Figure 1:**
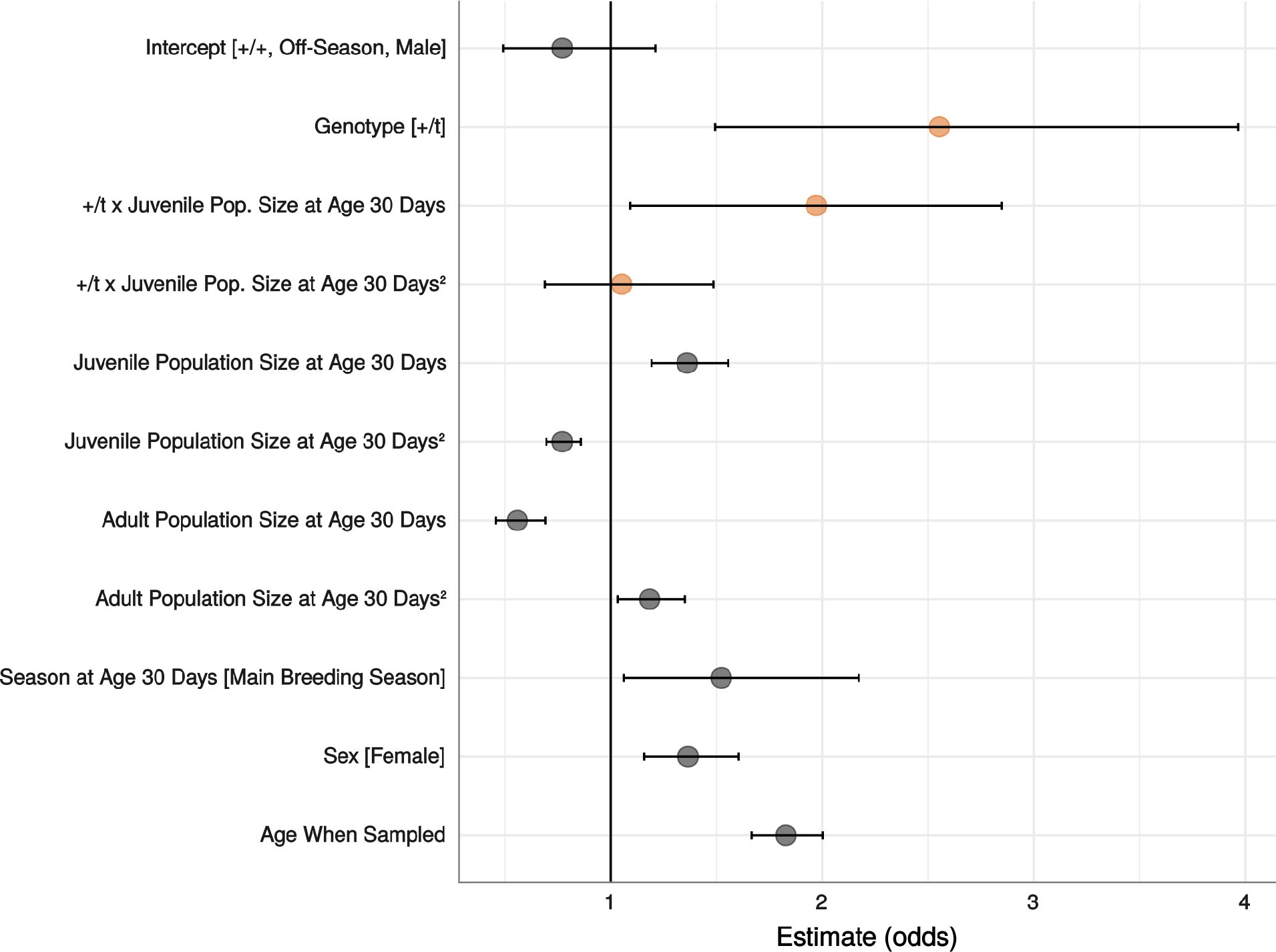
Effect estimates in odds with 95% confidence intervals of the most informative juvenile emigration model (*N* = 2938). The level of a categorical variable for which the effect is calculated is given in square brackets. Continuous variables are scaled. Interactions are indicated with an “x” between the variables. *t* main effect and interactions with *t* are highlighted in orange.

**Figure 2:**
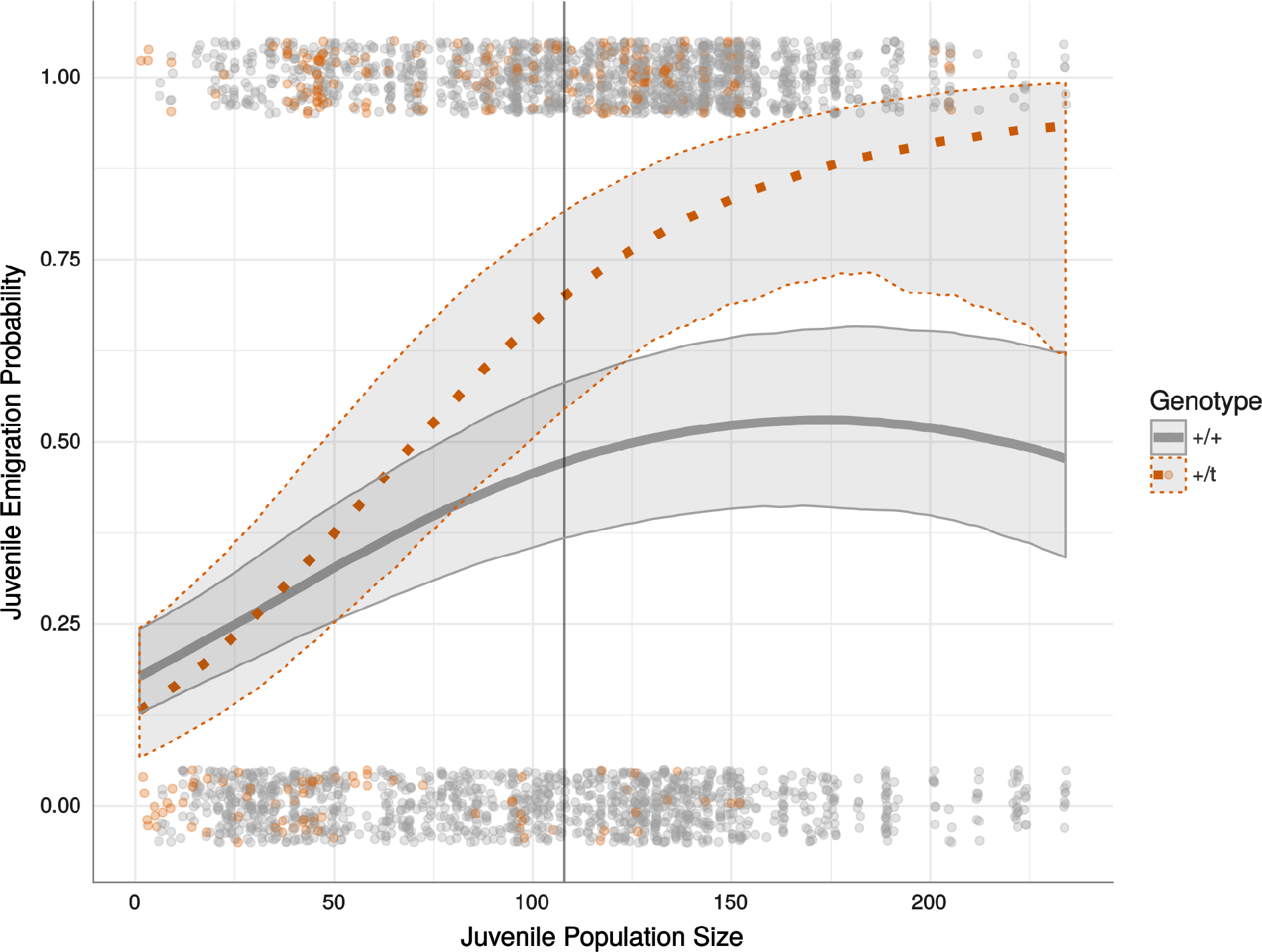
Predicted probabilities of juvenile emigration out of the study population (lines) with 95% confidence intervals and actual data points (top and bottom, jittered) of +/*t* (orange, dotted line) and +/+ (grey, solid line) individuals in varying juvenile population sizes (*N* = 2938). This exemplary plot is based on predictions from the most informative emigration model (emigration model 2) for a female born in the off-season in average adult population size for no specific birthyear (fixed effects only). The vertical line indicates the mean juvenile population size.

### Migration within the population

If +/*t* juveniles are more prone to emigrate from the population, they might also be more likely to migrate within this large population between sub-populations. To test this, we took the same dataset as in the emigration analysis, but restricted it to individuals that survived and remained in the population until adulthood (see Methods for details). Of the 901 individuals analysed, 11.1% migrated as juveniles within the population, i.e. they were found in a different sub-population as adults than they were last seen in as pups. 19.7% of the 61 +/*t* migrated within the population as juveniles compared to 10.5% of 840 +/+, a statistically significant difference (χ^2^ = 3.99, *df* =1, *p* = 0.046).

However, when controlling for other variables in a GLM, the results were less clear (Table 2). Adding the genotype to a null model with control variables did not improve the model, while a model that contained two genotype interactions, one with adult population size and one with sex was found to be significantly more informative than the null model. We used this model to visualise and estimate effect sizes. The predictor estimates of this model had large confidence intervals (see Figure 3). The estimate for +/*t* became negative when the interaction with adult population size was added to the model. The reason for that was that +/*t* juveniles were more likely to migrate within the population than +/+ only when the adult population size was small (see Figure 4). +/+ mice were least likely to migrate in mean juvenile population sizes and their probability slightly increased in both directions. A model with an interaction with juvenile population size instead of adult population size revealed a very similar interaction and adding both interactions did not improve the model any further, suggesting that population size *per se* is the relevant metric. Furthermore, male +/*t* were more likely to migrate than female +/*t*. Importantly, only few individuals were +/*t*, which decreased our power to detect and describe effects accurately. Mice born in the main breeding season were less likely to migrate within the population (Figure 3), but there was no difference between the genotypes (Table 2). Pup body mass was a positive predictor of migration, but did not differ between the genotypes or change the genotype’s effect (see migration models 10-12 in the S3 Table).

**Table 2:** Overview of models of juvenile within-population migration. Asterisks in model terms indicate interactions. Comparison shows against which other model the model in the row was evaluated. *LRT* indicates the likelihood ratio test statistic of the observed dataset. The *p*-value is the fraction of simulated datasets with *LRT* larger than the observed *LRT* (see Methods). Runs indicate the absolute values on which the *p* is based. The Δ*AIC* is given for comparison with other statistical approaches.

| Models | Formula | Comparison | $LRT$ | $p$ | Runs | $\Delta AIC$ |
| --- | --- | --- | --- | --- | --- | --- |
| Null model with covariates | ~ juvenile population size<br>+ juvenile population size<sup>2</sup><br>+ adult population size<br>+ adult population size<sup>2</sup><br>+ season + sex<br>+ age when sampled | NA | NA | NA | NA | NA |
| Model 1 | ~ genotype<br>+ null model variables | Null model | 0.21 | 0.65 | 6521/10000 | +1.79 |
| Model 2 | ~ genotype * juv. pop. size<br>+ genotype * juv. pop. size<sup>2</sup><br>+ model 1 variables | Model 1 | 5.66 | 0.09 | 876/10000 | -1.66 |
| Model 3 | ~ genotype * ad. pop. size<br>+ genotype * ad. pop. size<sup>2</sup><br>+ model 1 variables | Model 1 | 5.94 | 0.08 | 748/10000 | -1.94 |
| Model 4 | ~ genotype * juv. pop. size<br>+ genotype * juv. pop. size<sup>2</sup><br>+ model 3 variables | Model 3 | 3.34 | 0.26 | 2636/10000 | +0.66 |
| Model 5 | ~ genotype * season<br>+ model 1 variables | Model 1 | 6.47 | 0.13 | 1269/10000 | -0.47 |
| Model 6 | ~ genotype * sex<br>+ model 1 variables | Model 1 | 4.26 | 0.04 | 441/10000 | -2.26 |
| Model 6 | *as above* | Null model | 4.46 | 0.12 | 1207/10000 | -0.46 |
| Model 7 | ~ genotype * ad. pop. size<br>+ genotype * ad. pop. size<sup>2</sup><br>+ model 6 variables | Model 6 | 6.44 | 0.05 | 533/10000 | -2.45 |
| Model 7 | *as above* | Null model | 10.91 | 0.04 | 440/10000 | -2.91 |
| Model 8 | ~ genotype * juv. pop. size<br>+ genotype * juv. pop. size<sup>2</sup><br>+ model 6 variables | Model 6 | 6.38 | 0.07 | 680/10000 | -2.38 |
| Model 9 | ~ genotype * juv. pop. size<br>+ genotype * juv. pop. size<sup>2</sup><br>+ model 7 variables | Model 7 | 3.16 | 0.31 | 3124/9996 | +0.84 |

**Figure 3:**
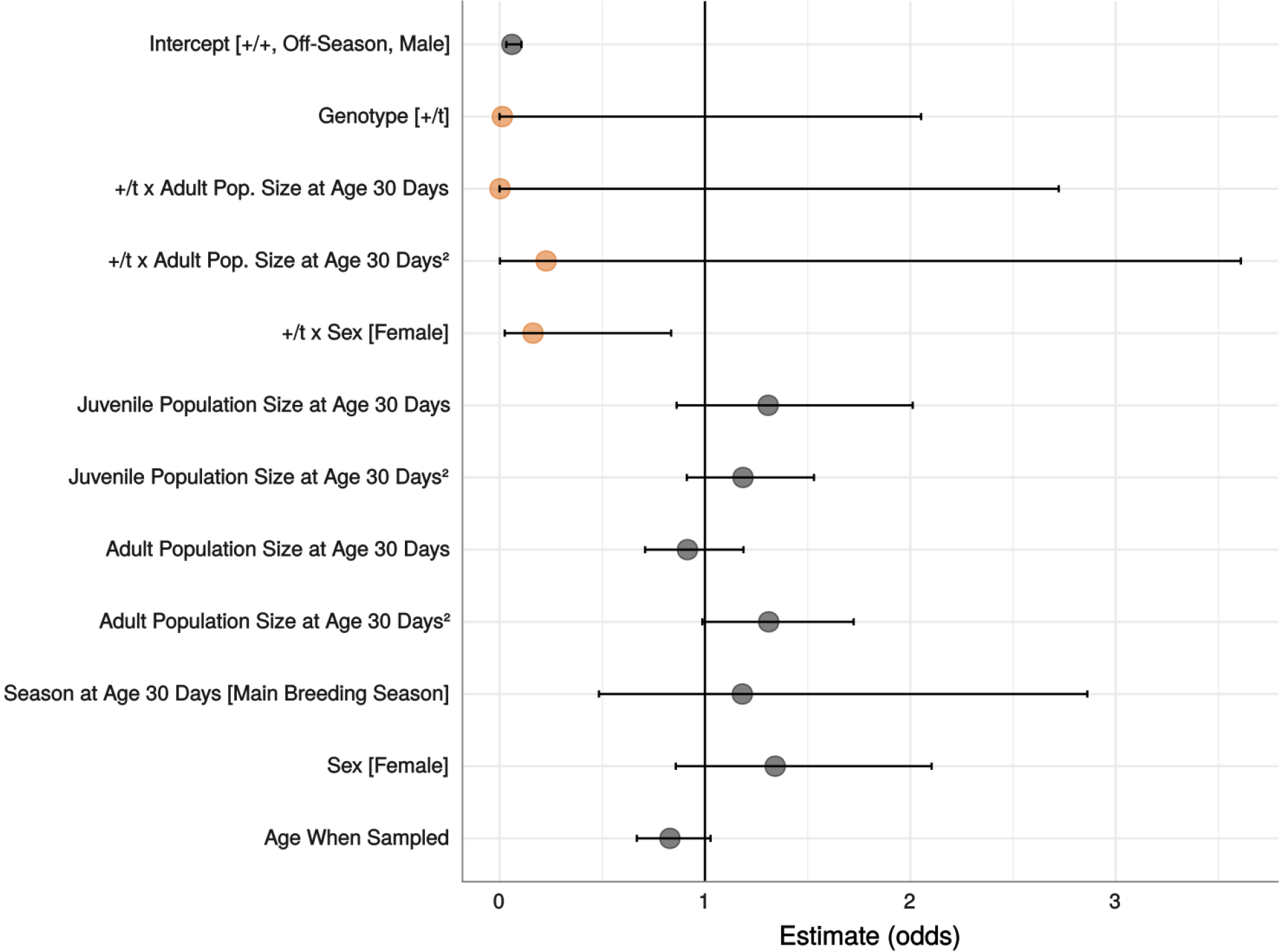
Effect estimates in odds with 95% confidence intervals of the most informative juvenile within-population migration model (*N* = 901). The level of a categorical variable for which the effect is calculated is given in square brackets. Continuous variables are scaled. Interactions are indicated with an “x” between the variables. *t* main effect and interactions with *t* are highlighted in orange.

**Figure 4:**
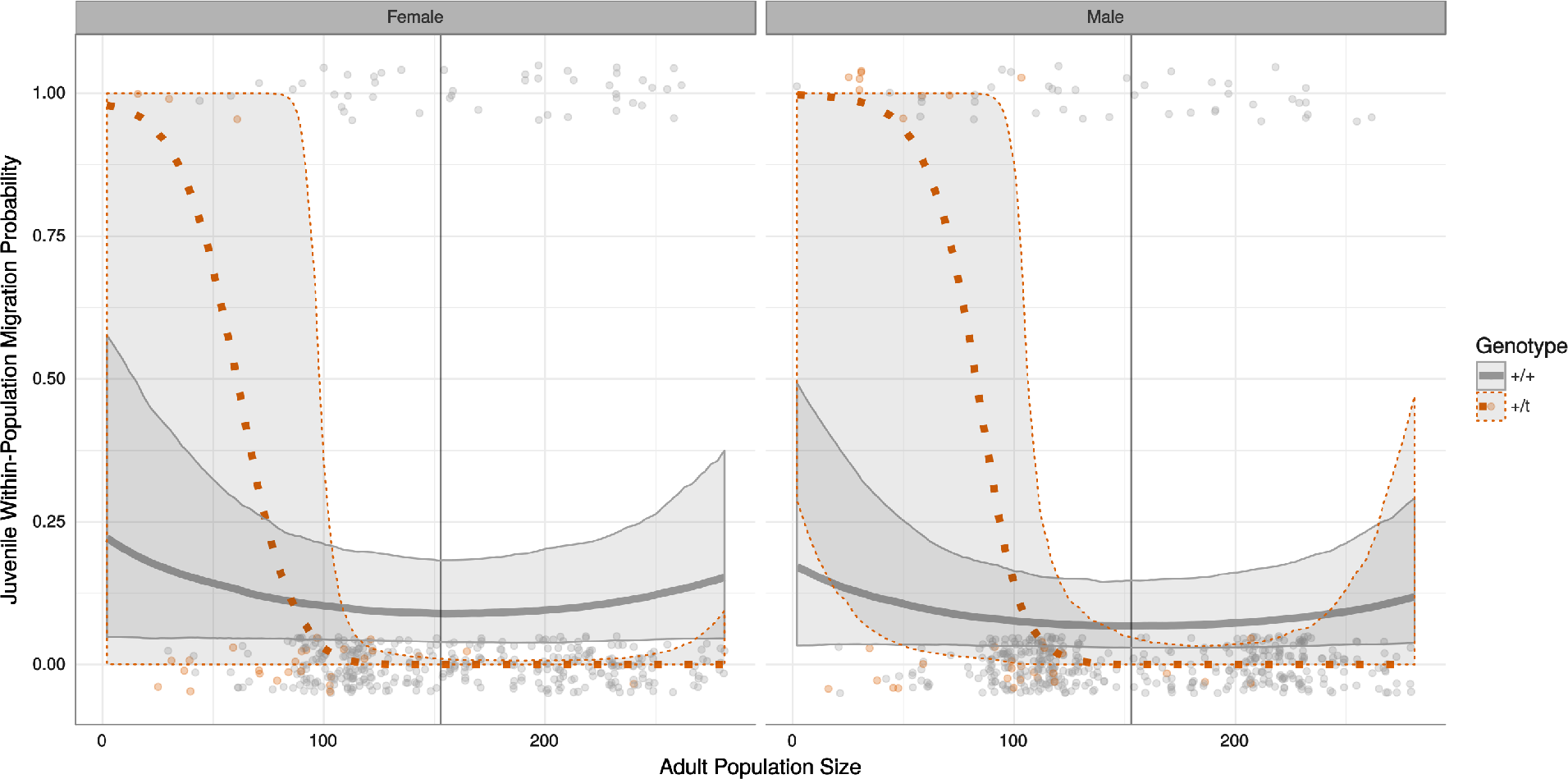
Predicted probabilities of juvenile within-population migration (lines) with 95% confidence intervals and actual data points (top and bottom, jittered) of +/*t* (orange, dotted line) and +/+ (grey, solid line) individuals in varying adult population sizes, separated by sex (*N* = 901). These plots are based on predictions from the most informative within-population migration model (migration model 7) for individuals that were born in the off-season in average juvenile population size. The vertical line indicates the mean adult population size.

## Discussion

We provide evidence for a higher migration propensity of +/*t* juveniles compared to +/+ juveniles. We found that carrying the *t* haplotype is a strong positive predictor for juvenile emigration out of our study population. Our hypothesis that +/*t* should be selected to increase migration propensity was also modestly supported by a +/*t* bias in migratory movements within the population. Given that variation in behaviours related to dispersal is generally heritable to moderate degrees [45], a manipulation by the *t* in the *t*’s favour is a probable explanation. Our results further suggest that the *t* increases emigration propensity particularly in denser populations. This is plausible because the *t* should be less fit in denser populations due to an increase in sperm competition [20,30]. The +/*t* that did not emigrate from the population were found to be more likely to migrate within the population when adult densities were low. A possible explanation for this could be that there was more open habitat available when less adults were in the population and the migration-prone +/*t* were able to migrate within the population instead of needing to leave it.

We did not find a different effect of sex between the genotypes in our emigration analysis, but did find one in the within-population migration analysis. The lack of difference agrees with a theoretical model that showed that *t* migration propensity manipulation need not be biased towards males (in which *t* drives), because migration of both male and female +/*t* was found to be more effective than male-only migration [38]. However, +/*t* males were more likely than females to migrate within the population as juveniles. The test of this interaction was exploratory and not driven by a hypothesis. The result may reflect sex-specific costs and benefits of within-population migration for +/*t* mice. It is interesting, but needs further verification, particularly given that the emigration analysis with its larger dataset does not show this pattern.

One drawback of our emigration analysis is that it is based on an indirect measure of emigration, which we expect to be less precise. Despite that, we detected a strong signal. We considered alternative explanations of the strong +/*t* emigration bias. We tested for a difference in juvenile mortality, but did not find one, which is further supported by a lack of difference in pup survival until weaning from lab-bred mice taken from the same population [26]. We found a slightly increased pup body mass for +/*t*, but showed that this was unlikely to drive the emigration difference. Furthermore, there is evidence from another lab study that +/*t* and +/+ from the same study population do not differ in adult body mass (males and females) [17]. Differences in social dominance could be another explanation for emigration patterns. Studies looking at dominance either found less dominant +/*t* males [46], more dominant +/*t* males [47], or no difference in dominance between males and less dominant +/*t* females [35,48]. However, if dominance differences were the cause of our emigration results, we might expect to see an informative interaction between sex and genotype, unless our population is one where both +/*t* males and females are less dominant. Furthermore, we know from previous analyses that +/*t* males do not differ in survival from +/+ and +/*t* females live longer than +/+ in our population [20]. Survival can predict dominance in house mice [48] and thus there is no clear evidence that dominance differs between the genotypes in our population and then drives emigration patterns.

Generally, the increased migration propensity of +/*t* could help to explain why the *t* continues to exist in nature despite its homozygous and heterozygous fitness costs due to recessive lethals and low sperm competitiveness. Compared to a *t* variant that does not influence migration, variants of the *t* that increase migration propensity could have an increased chance of reaching or founding populations where there are few other +/*t* and polyandrous matings are less frequent. The *t* is expected to rapidly increase in frequency given such circumstances [20,31,49–51]. We expect a *t* variant that manipulates migration propensity to increase the odds that its carrier is in an environment that allows for rapid increases in *t* frequency. Thus, it would likely out-compete *t* variants that did not affect migration (and these could go extinct locally). Competition between *t* variants is consistent with genetic evidence that a single *t* haplotype variant recently replaced previous variants in a sweep [52]. We do not know how an increased migration propensity could be encoded within the *t* haplotype, but the *t* comprises several hundred genes that are protected from recombination [25]. Alternatively, instead of manipulation by the *t*, the increased migration propensity could also be a response by the rest of the genome to the presence of the *t*, if increasing migration propensity is increasing the fitness of the rest of the genome when *t* is present. More work is needed to better understand this interesting dynamic.

Emigration is only the first step of successful dispersal. Emigrants also need to breed as an immigrant or founder, which is challenging for mice [53]. Unfortunately, there were too few +/*t* that migrated within the population for us to analyse their breeding success. However, Anderson et al.[54] were able to “infect” an island population with the *t* haplotype by manually migrating +/*t*. Although the *t* was able to establish itself in the initial area over a period of a few years, it did not spread much across the island. For Pennycuik et al.[55], introducing the *t* to an enclosure was more difficult. However, they managed to do so when there were open territories in the population. They also reported many of the +/*t* males and females migrating between sub-populations. However, the *t* was almost extinct two years later, at the end of the study. It is evident from these experiments that there will be many populations to which the *t* cannot disperse successfully. In our study population we have no evidence for immigration of any +/*t* individuals (unpublished). This makes increased migration propensity counter-intuitive because the migration will often fail. Still, because not migrating is also not beneficial for the *t*, it makes migration attempts potentially even more necessary for the *t*’s fitness.

When house mice invade an island that has evolved without mammalian predators, their presence can be very damaging to the ecosystem [56–58]. Recently, efforts are being made to use a modified *t* haplotype for potential eradication of such house mouse populations [44,59,60]. The *t*_*SRY*_ variant is a *t* haplotype that is synthetically combined with the male-determining gene *SRY*. Every +/*t*_*SRY*_ individual is thus male. Due to the *t*’s transmission advantage, more than 90% of the offspring of a +/*t*_*SRY*_ are then male, which could then drive populations extinct via lack of one sex [43,61,62]. So far, only some of the *t*’s characteristics have been explicitly considered in trying to facilitate the use of *t*_*SRY*_ to eradicate wild populations [43]. However, accounting for the entirety of the known attributes of the *t* is crucial to successfully predict how a a synthesized variant works in the field, in particular the *t*’s sperm competition disadvantage [17]. Increased migration propensity would likely aid in the distribution of +/*t*_*SRY*_ mice to target locations, but could also increase the possibility of *t*_*SRY*_ reaching populations it was not intended for.

To our knowledge, this is the first evidence of manipulation of migration propensity by a selfish genetic element. Our results should be of broad interest. First, they have implications for research on other selfish genetic elements, considering low sperm competitiveness is expected in many male meiotic driver systems like the *t* [14–16,18,63]. Recessive deleterious alleles and therefore frequency-dependant fitness would also be expected in other meiotic drivers, because without negative fitness effects the driver would spread to fixation [64,65].

This would provide further advantages for migratory variants of these drivers. Similarly, parasites could also benefit from manipulating dispersal behaviour [40]. Second, the recent work on artificial gene drive systems based on the *t* haplotype will benefit from incorporating as many traits of the *t* as are available. A difference in migration propensity could have important implications for such a system. Third, a selfish genetic element affecting migration propensity could be an important finding for research on dispersal and migration in general. Dispersal attempts are risky [66] and the different selective pressures for the *t* and similar elements could help to explain better when this behaviour - that often results in no fitness gains - is most beneficial. Therefore, arms races like the one studied here could be a causal mechanism driving the evolution of dispersal. We will further investigate this new direction in *t* haplotype research with theoretical and experimental approaches.

## Methods

### The population

We analysed data that were collected between the years 2004 and 2012 in a free-living house mouse *Mus musculus domesticus* population in an old barn near Zurich, Switzerland [67] under permits 26/2002, 210/2003, 215/2006, 51/2010 from the Swiss Animal Experimentation Commission. As house mice in Switzerland live commensally with humans, we provided a human-made and provisioned environment similar to that found in barns housing animals, but easier to monitor. We provided food (a 1:1 mix of rolled oats and hamster food from Landi AG, Switzerland) and water regularly *ad libitum*. The barn is divided into four similarly sized sectors [67]. However, mice can easily travel between these sectors by passing through holes in the walls or climbing over them. The mice can also freely enter and leave the population. This migration could not be monitored directly due to the numerous and unpredictable exit routes that mice use (that were however small enough to exclude predators). We used a indirect measure of emigration (see “Definition of emigration”). We considered individuals from 1 to 16 days as pups, then (when they begin to be weaned) as juveniles before reaching 17.5 grams in body mass, which is when we classified them as adults, as females do not breed until they exceed this body mass [32]. The sex ratio of the population was roughly equal (48% of the individuals in this analysis are female).

### Monitoring

When pups reached 13 days of age (allowing for ±2 days of difference from this), they were ear-punched to provide a DNA sample. Every 10 to 13 days, the barn was searched for new litters. The age estimation was based on the developmental state of the pup [32]. Every 7 weeks, on average, every individual in the barn was caught. On this occasion, all individuals above 17.5 grams in body mass received an RFID transponder and were then considered adults. On average in the years studied, 16.1% of the population received a transponder (was newly classified as an adult) on such a capture event. Additionally, we regularly searched the barn visually and with transponder scanners for dead individuals or lost transponders. When found, dead individuals were removed and identified via their transponder or a new genetic sample. Finally, there is an automatic antenna system since 2007 in our population that tracks exits and entries of transpondered mice into and out of our 40 nest boxes [67]. We used these data in addition to data from manual checks to determine when an adult individual was last detected in the population if it was never found dead. This was relevant for the population size calculations, see “Controlling variables”.

### Identification

We genetically identify each individual as a pup, as a newly classified adult, or as a corpse if found dead without transponder. We do so based on multi-locus genotypes based on 25 micro-satellite loci [68]. The genotypes allow us to link individuals as pups to their adult transponder ID or to a corpse, allowing for one allelic mismatch using the software CERVUS [69]. We use the micro-satellite locus *Hba-ps4* that has a 16-bp *t* specific insertion [70] to identify the *t* haplotype. Sexing of individuals was performed by testing for the presence of Y-chromosome-specific micro-satellite markers Y8, Y12, and Y21[71].

## Definition of emigration

### Emigration out of the population

Individuals that fulfilled all of the following criteria were classified as juvenile emigrants: 1) Individual was genotyped as a 13± 2 day old pup, 2) its genotype never matched to an adult’s sample, and 3) also never to a corpse’s sample. Following this definition, the time at which the individual could have emigrated must have been between 13 ± 2 days of age and an adult age (defined by body mass as described earlier) and therefore the individual was a juvenile. Consequently, individuals that left the barn as adults are not classified as emigrants in this analysis, but are instead treated as juvenile non-emigrants. We excluded individuals born in the year 2005 from the analysis because monitoring was considerably less intense in this year and thus there is a larger potential to misclassify individuals that died within the population as ones that emigrated. Therefore, we analyzed 7 birthyears (2004 & 2006-2011) in which the *t* was present in the population (it then went extinct). We also excluded individuals (213) about whom we did not have enough information (such as incomplete genotype or conflicting sex information) from the analysis. Furthermore, we removed those that died as juveniles, because we cannot know whether they might have emigrated later (218). The absolute numbers provided in brackets are sequential decreases in sample size, e.g. the 218 dead juveniles were excluded after the other dataset restrictions were made.

### Migration within the population

We defined the four distinct sectors within the population described earlier as sub-populations between which mice can migrate. We did so for two main reasons: 1) From earlier analyses [72] and anecdotal observations we know that the dividing walls between the four sectors are social but not physical barriers for the mice. While mice are regularly seen moving within each sector, movements and social interactions between the sectors are less frequent. [72] 2) When considering the location of adults during our regular monitoring of the population (the data used here), 61% of adults (in their adult lifetime) that were found at least 9 times were found within the same sector every time. 31% were found in two sectors in total, 7% in three, and less than 1% in all four. We defined juvenile within-population migrants as individuals that were first found as adults in a different sector than they were last seen in as pups. Thus, these individuals migrated at an age where they were older than a 13 ± 2 day old pup (when we sampled them the first time), but younger than when they would reach adult body mass. We therefore expect both migratory behaviours analysed in this study (within and out of the population) to have taken place at similar ages. The dataset was based on the same restrictions made for the emigration analysis, except that only those individuals that stayed in the population until adulthood could be analysed.

### Controlling variables

We estimated the population size at any date by counting all individuals that were alive. In cases where an individual left the population or died but was not found, we used the date an individual was last detected in the barn as the last date present in the population. This date is based on both manually locating (in regular population monitoring) the animal or information from our automatic antenna system. Furthermore, a large proportion of the individuals disappeared from our population before they receive their RFID transponder (the emigrants in this paper). Individuals that disappeared in this way were counted for 30 days from the time of their birth on as part of the population. This cut-off is based on a handful of individuals that reached the body mass we designate as minimum for the transponder (17.5 g) at 35 days of age, reports of an early dispersal phase in 30 day old juveniles [73], and a weaning age (nutritional independence and end of active maternal care) in mice of about 23 days [74,75]. Therefore, it is a conservative estimate of the minimum amount of time an emigrant would spend in the population after birth. However, the results of this study do not change fundamentally when this time frame is increased (we used 50 days of age as an alternative cut-off, see S1 Table).

We subdivided the population size into adult and juvenile population size. We did so because the two could influence the mice differently. We do not know how individual mice decide whether they emigrate, therefore we wanted to disentangle the two variables that could reflect the current and the future reproductive environment. The two population sizes are correlated, but do not explain much of variation of each other (linear model with *R*^2^ = 0.05). For the purpose of consistency within this analysis, individuals that remained in the population until adulthood were counted from age 31 days on as part of the adult population (and before as juveniles and pups), whereas individuals that were never considered as adults were only counted for 30 days as juveniles and never as adults.

We defined the months April to September as the main breeding season, because these are the 6 months with the highest counts of new pups. The remaining months (October to March) were defined as the off-season. 87% of the birth dates in our dataset fall within the main breeding season. Because of a possible immeasurable multitude of inter-annual variation in the environment (like temperature or noises in the area) that could possibly affect migration propensity, we controlled for the the year of birth (*N* = 7) as a random effect in the emigration models. Finally, we also controlled for the age when individuals were first sampled (between 11 and 15 days of age with most being sampled at 13 days). We did so because preliminary data visualisations revealed a relation between this age and emigration.

## Statistical analyses

### Emigration out of the population

To test whether the relation between genotype and emigration is statistically significant when controlling for other factors, we used a generalized mixed effect model with a binomial distribution and a logit-link function and fit by maximum likelihood. All statistical analyses and figures were done in *R* 3.4.2 [76] with *RStudio* [77] and the packages *ggplot* 2 2.2.1 [78], and *lme4* 1.1-14 [79]. The function *glmer* of the package *lme4* was used for the modelling. The dependent variable is a binary variable (1 when the individual emigrated as a juvenile and 0 if it did not). The independent variables are adult and juvenile population size (each scaled by the standard deviation, centred at zero, and fitted as linear and quadratic terms to estimate slope and shape of the prediction), the season (off-season as intercept), the individual’s sex (male as intercept), and its genotype (+/+ as intercept). The population sizes and the season are taken from 30 days after an individual’s birth to reflect the environment that the juvenile was facing around the time when it either did or did not emigrate. The year of birth is used as a random effect. We used the function *confint* of package *lme4* for parameter confidence intervals in Figure 1 with the built in basic bootstrapping method and 1000 simulations. To estimate the prediction and confidence intervals of the fixed effects of the most informative model, we used the function *predictInterval* of *merTools* 0.3.0 [80] with its integrated bootstrapping method with 1000 simulations, using the median and a confidence interval of 95% for Figure 2.

We used the R package *pbkrtest* 0.4-7 [81] for parametric bootstrapping based model comparisons. Each dataset was simulated 10000 times. The *p*-value is based on the *PB* statistic provided by the function *PBmodcomp*. It represents the fraction of likelihood ratio test (*LRT*) values of the simulated (bootstrapped) datasets that were larger or equal to the observed *LRT* value. Some of the runs can result in negative values of the *LRT* statistic. These runs are excluded automatically and the number of used runs along with the number of runs where the *LRT* is more “extreme” than the observed *LRT* are provided in the Results. We used a significance level of 5%. With this, we test the significance of 1. the genotype’s effect, 2. the interaction between genotype and juvenile population size, and 3. the interaction between genotype and adult population size. 1. was tested by comparing a model with all the controlling variables (“emigration null model”) but without genotype as a predictor with a model that had the same set of predictors plus the genotype. 2. and 3. were each tested by comparing a model with all controlling variables and the genotype (“emigration model 1”) to a model that also included the interaction (“emigration model 2” with an interaction with juvenile population size and “emigration model 3” with adult population size). We also tested whether the interactions become more informative if both interactions (with juvenile *and* adult population size) are in the model (“emigration model 4”). We also list Δ*AIC* values in the results to ease understanding, but do not use them for interpretation. The full output of each of these models can be found in the SI. Lastly, we tested the interaction of genotype and season (“emigration model 5”) as well as sex (“emigration model 6”) to explore relationships that we did not hypothesize. We did so by comparing the most informative model with and without an interaction of genotype and sex or season.

To test whether pup condition differences could be an alternative explanation for the emigration differences, we used the same environmental variables to conduct a linear mixed model that predicts pup body mass and then compared this model to one that also included the genotype as an effect (“body mass null model” & “body mass model 1” in the SI). We then added pup body mass as a predictor to our emigration null model (“emigration model 7”, SI) and our most informative emigration model (“emigration model 8”, SI) to test whether a) emigration is predicted by pup body mass and b) the genotype explains the same variation as does the pup body mass. All analyses that included body mass are reduced in their sample size by 40 individuals for whom we did not have this information.

### Migration within the population

For this analysis, we have a reduced sample size because only mice that stayed alive and remained within the population until adulthood can be analyzed. We also excluded one more birthyear because in 2011 no +/*t* stayed in the population until adulthood. We analyzed 901 mice. The number of +/*t* in this dataset is small (61), which complicates statistical analyses. We first compared the numbers of juvenile migrants between the genotypes with Yates’s χ^2^ test using R. Then, without subdividing the data into years of birth (i.e. no random effect), we used a generalized linear model to control for the same variables as in the emigration analysis. We selected the most informative model using the same parametric bootstrapping approach as in the emigration analysis. We used the *confint* function of package *MASS* 7.3-47 [82] to estimate the confidence intervals in Figure 3 using a 95% confidence interval. For the prediction and its 95% confidence interval in Figure 4 we used the R base function *sample* to draw from 1000 bootstrapped simulations.

## Acknowledgements

We are particularly grateful to Barbara König for her perseverance in keeping this long term field study going and for her generous support, and to her and all others who have contributed to collection of the data. We also thank Jari Garbely for genetic lab work. Lastly, we thank Barbara König, Tom Price, Erik Postma, and Andri Manser for comments on an earlier version of this manuscript. This study was funded by the SNF (31003A-120444, 310030M_138389, 31003A_160328), the University of Zurich, the Promotor Foundation, Julius Klaus Foundation, and the Claraz-Stiftung.

## Author contributions

The study was conceived and the manuscript written by JR and AL. The data were collected and the genetic analyses performed by AL and her team. Statistical analyses were performed by JR.

---

**S1 Table. Full model outputs of the juvenile emigration models.**

**S2 Table. Full model outputs and comparison between the pup body mass models.**

**S3 Table. Full model outputs of the juvenile migration models.**

